# The habenula clock influences prediction of danger

**DOI:** 10.1101/2020.04.29.067108

**Authors:** Adriana Basnakova, Ruey-Kuang Cheng, Joanne Chia Shu Ming, Giuseppe D’Agostino, Suryadi, Germaine Tan Jia Hui, Sarah R. Langley, Suresh Jesuthasan

**Affiliations:** School of Biological Sciences, University of Manchester, UK; Institute of Molecular and Cell Biology, Singapore; Lee Kong Chian School of Medicine, Nanyang Technological University, 59 Nanyang Drive, Singapore 636921.; School of Physical and Mathematical Sciences, Nanyang Technological University Singapore.

**Keywords:** anxiety, circadian clock, habenula, spontaneous activity, predictive coding

## Abstract

The response of an animal to a sensory stimulus depends on the nature of the stimulus and on predictions mediated by spontaneous activity in neurons. Here, we ask how circadian variation in the prediction of danger, and thus the response to a potential threat, is controlled. We focus on the habenula, a mediator of threat response that functions by regulating neuromodulator release, and use zebrafish as the experimental system. Single cell transcriptomics indicates that multiple clock genes are expressed throughout the habenula, while quantitative in situ hybridization confirms that the clock oscillates. Two-photon calcium imaging indicates a circadian change in spontaneous activity of habenula neurons. To assess the role of this clock, a truncated *clocka* gene was specifically expressed in the habenula. This partially inhibited the clock, as shown by changes in *per3* expression as well as altered day-night variation in dopamine, serotonin and acetylcholine levels. Behaviourally, anxiety-like responses evoked by an alarm pheromone were reduced. Circadian effects of the pheromone were disrupted, such that responses in the day resembled those at night. Behaviours that are regulated by the pineal clock and not triggered by stressors were unaffected. These findings establish that the habenula clock regulates the expectation of danger, thus identifying one mechanism for circadian change in the response to a stressor.

## 1. Introduction

Expectations play a major role in determining the response of a person or animal to a sensory stimulus or situation. In this framework, known as predictive coding, the brain is not a passive recipient of stimuli, but actively generates predictions that are updated by experience (Barrett, 2017; Friston, 2018). Thus, encountering a cue such as the smell or sound of a predator increases the expectation of danger, turning a normally innocuous stimulus into a threat. Abnormal expectations, or predictions, have been proposed to underlie stress-related conditions such as post-traumatic stress disorder. Predictions are mediated by spontaneous activity in the brain (Berkes et al., 2011), while updating of expectations is enabled by the release of neuromodulators. Dopamine and serotonin contribute to the comparison of current input with pre-existing expectations. When outcomes surpass predictions, dopamine release by neurons in the midbrain increases; when there is disappointment, dopamine release is inhibited (Schultz, 1998). Serotonin release by the raphe provides an indication of the magnitude of the prediction error (Matias et al., 2017). Acetylcholine appears to have a role in setting the precision of predictions (Moran et al., 2013). Behaviour and emotional states can thus be understood to be the result of genetically encoded programs that are continuously modified by predictions, enabling the animal to respond optimally to perceived reality.

The habenula, which contains several subdomains, has emerged in recent years as an important regulator of several neuromodulators involved in predictive coding. The medial habenula in zebrafish (Hong et al., 2013), like that of mouse (Ren et al., 2011), includes cholinergic neurons. Disruption of specific subdomains of the medial habenula leads to anxiety-like behaviour (Agetsuma et al., 2010; Lee et al., 2010) or aversion (Hsu et al., 2014) that is experience dependent. The lateral habenula regulates midbrain dopaminergic and serotonergic neurons, and phasic activity occurs here when there is an unexpected punishment or absence of reward (Matsumoto and Hikosaka, 2009). As an animal learns the actions that prevent punishment, this activity lessens (Amo et al., 2014). In addition to activity that is evoked by sensory stimuli or perceptions of reward, the habenula displays spontaneous activity (Jetti et al., 2014; Kim and Chang, 2005). Excessive levels of burst firing, which can be interpreted as constant negative prediction, has been detected in the lateral habenula of animals with depressive behavior (Andalman et al., 2019; Yang et al., 2018). To understand how mood and behaviour are shaped, it is thus important to not only understand how the habenula responds to reward and sensory input, but to also investigate how spontaneous activity is regulated.

One factor that affects the level of spontaneous activity in neurons is the molecular clock. The molecular clock in most organisms consists of a transcriptional-translational feedback loop with a period of ~ 24 hours (Takahashi, 2017). Core components of the clock in animals include heterodimers of Clock and Arntl (Bmal), which enter the nucleus in the daytime. Here, they drive transcription of *period* (*per*) and *cryptochrome* (*cry*) genes, which then inhibit the Clock-Arntl heterodimer. Clock and Arntl also control transcription of channels that affect resting membrane potential in neurons, driving this close to threshold and thus increasing spontaneous firing rate in the daytime (Harvey et al., 2020). Components of the clock machinery are found in nearly all cells, including those of the habenula (Baño-Otálora and Piggins, 2017; Mendoza, 2017), and the habenula molecular clock continues cycling for multiple days even in the absence of the main pacemaker of the brain, the suprachiasmatic nucleus (Guilding et al., 2010; Salaberry et al., 2019). Spontaneous firing in the habenula of rat (Zhao and Rusak, 2005) and mouse (Guilding et al., 2010; Sakhi et al., 2014a, 2014b) varies across a circadian cycle, in both the lateral and medial habenula. The significance of the habenula clock is unclear, however. Given the role of habenula activity in predictive coding, we hypothesize that the clock causes a circadian change in behaviours that are modified by recent experience. To test this, we use the zebrafish as the experimental system. We examine anxiety-like behaviour, which is increased by expectation of danger.

## 2. Materials and Methods

### 2.1 Single cell data reanalysis

Count tables were downloaded from Gene Expression Omnibus, accession number GSE105115. All processing downstream was done using R/BioConductor (R 4.0.2/Bioconductor 3.12) packages designed to handle single cell data using the SingleCellExperiment class (v. 1.12.0) (Amezquita et al., 2020). A detailed explanation with code and figures is provided as a GitHub repository at https://github.com/langleylab/habenula_reanalysis. Briefly, gene-level count tables from the SmartSeq2 experiment were merged retaining the information on the plate of origin. Genes that were not detected in any barcode were removed. Barcodes were discarded according to the following criteria: low expression, low number of genes detected, high percentage of *malat1* expression, high percentage of mitochondrial transcripts expression and high percentage of ribosomal transcript expression. Thresholds for every criteria were set as 3 Mean Absolute Deviations (MADs), with the exception of mitochondrial % which was set at 4 MADs. Outlier calculations were done separately in each plate using the *isOutlier* (using the *batch* argument) and *perCellQCMetrics* functions from the *scater* package (v. 1.18.6) (McCarthy et al., 2017). These filters yielded 17673 genes and 1140 barcodes for downstream analyses. Barcodes were then clustered using the *quickCluster* function from the *scran* package (v. 1.18.5 (Lun et al., 2016)), blocking for the plate of origin. Per-cell size factors were then determined by deconvolution using *scran*, and they were used to perform multi-batch normalization (batching by plate) using the *multiBatchNorm* function from the *batchelor* package (v. 1.6.2 (Haghverdi et al., 2018)). Highly variable genes were estimated on the multi-batch normalized data blocking by plate using the *modelGeneVar* function from the *scran* package, which estimates the mean-variance trend and decomposes variance for each gene in technical and biological components according to the deviation from the trend. Variances were combined across plates using the *combineVar* function from *scran*, and the 2000 genes with the highest biological variance were selected using the *getTopHVGs* function from *scran*. These were used as an input to perform dimensionality reduction via Principal Component Analysis as implemented in the *BiocSingular* package (v. 1.4.0, (Lun, 2020a)).

The first 50 principal components were then used as an input to perform batch (plate) effect correction via the *fastMNN* function from *batchelor*, which determines a correction vector in PCA space using mutual nearest neighbour pairs to estimate the amount and direction of necessary correction. The dimensionality of the corrected space, i.e. the number of most relevant dimensions for subsequent steps, was determined using the *maxLikGlobalDimEst* function from the *intrinsicDimension* package (v. 1.2.0 (Johnsson et al., 2014)). The value for the *k* parameter for this function (k = 30) was determined as the rounded square root of the number of cells, and the estimation yielded 13 dimensions. Subsequently, the first 13 dimensions of the corrected space were used to run the t-stochastic neighbour embedding (t-SNE) algorithm for data visualization, and to construct a Shared Nearest Neighbour (SNN) graphs for clustering. Several SNN graphs were constructed at different resolutions, i.e. 10, 15, 30, 40 and 50 neighbours were considered for the graph construction; all edges were weighted by the Jaccard similarity between the neighbours of each node. SNN graph construction was performed using the *buildSNNGraph* function from *scran*. Then, the Louvain community detection algorithm (as implemented in the *igraph* package (v. 1.2.6 (Csardi and Nepusz, 2006)) was used to identify clusters. Relationships between different clustering results (obtained by using different neighbours in the SNN graph construction) were inspected using a clustering tree drawn using *clustree* (v. 0.4.3 (Zappia and Oshlack, 2018)). Clusterings with 10 and 15 neighbours were further explored, as they yielded numbers of clusters similar to the ones in the original publication. For both 10 and 15 neighbours, cluster separation was evaluated using three methods: approximate silhouette width, neighborhood purity and pairwise modularity, as implemented in the *bluster* package (v. 1.0.0 (Lun, 2020b)) in the *approxSilhouette*, *neighborPurity* and *pairwiseModularity* functions. The 15 neighbour clustering showed slightly better separation so it was considered for downstream analyses. In order to reconstruct the original cell type identities as identified by the authors, the *AUCell* package (v. 1.12.0, (Aibar et al., 2017)) was used in conjunction with the cluster marker genes as published by the authors, although these marker genes were identified in the 10X dataset. Raw counts were ranked in each cell and the AUC for each marker set was calculated; each cell was assigned the identity yielding the highest AUC value using the 30% top ranking genes. AUCell is agnostic to the transcriptional space and to other intrinsic properties of the dataset, which will be influenced by differences in filtering and processing steps. Therefore the performance of AUCell was evaluated in this dataset calculating the approximate silhouette width of the cell type assignment, and it was compared to our clustering results by calculating pairwise Jaccard indices. Most identities (Hb1, Hb2, Hb5, Hb6, Hb9, Hb1, Hb14, Hb15 and olfactory neurons) are reasonably well separated, but only 4 (olfactory, Hb06, Hb03, and Hb08), are reproduced in our clustering with a Jaccard index above 0.5 with any single cluster. This is in line with what the authors of the original paper state regarding the poor performance of SmartSeq2 in terms of cell type recovery, and is also to be expected when using cell type markers identified in the 10X dataset.

For the 10X datasets (larval) no filtering was performed as the distribution of QC metrics was well within standard ranges, suggesting data were already filtered. Normalization, HVG estimation and dimensionality reduction were performed in the same way as described above, with the exception that there was no batch effect to account for and correct. An additional doublet detection step was included using the *scDblFinder* package (v. 1.4.0, PMID (Germain et al., 2020)), which flagged 172 barcodes as doublets and 4193 as singlets. Doublets were not removed at this stage of the analysis. The intrinsic dimension estimation (k = 66) yielded 12 dimensions to perform SNN graph clustering, which was performed using 5, 10, 15, 20, and 30 neighbours. In parallel, 36 dimensions were also considered as done by the authors of the original publication. For the 12-dimensional space, 5 and 10 neighbours were considered as they yielded numbers of clusters similar to those reported in the publication; for the 36-dimensional space, 10 and 15 neighbours were considered. However, in both 12 and 36 dimensions a cluster was entirely made up of doublets, so doublets were removed and the whole process (normalization, HVG selection and dimensionality reduction) was repeated. Cluster separation metrics (approximate Silhouette and modularity) favoured clustering with 10 neighbours (12 dimension) and 15 neighbours (36 dimensions). To reproduce the original clustering results the AUCell package was used, and Jaccard indices were used to match clusters to labels. In order to improve the label assignment, cells whose top AUC label showed less than a 20% difference from the second best label were classified as “ambiguous”. Ambiguous labels were then reassigned using the closest centroid in the approximate silhouette width calculation, thus making the labels conform more to the transcriptional space. Improvements in the reassignment of these labels were measured by comparing silhouette widths and the distribution maximum Jaccard indices for every cluster-AUC label pair before and after reassignment. Both metrics showed an improvement in both silhouette width and median best Jaccard index upon reassignment.

Adult zebrafish data was processed in the same above, accounting for batch effects (2 different captures). SNN graph construction with 4 neighbours in 16 dimensions was chosen following the same pipeline and reasoning used in the larval dataset. AUCell was used to determine identities using the markers provided by the authors of the original publication, and the reassignment procedure described above was used to correct ambiguous labels.

### 2.2 Transgenic lines

*TgBAC(gng8:GAL4)^c426^* (Hong et al., 2013) orT^*s1011t*^ (Scott and Baier, 2009) was used to drive expression in the habenula. Calcium imaging was performed using *Tg(elavl3:GCaMP6f)^a12200^* and *Tg(UAS:GCaMP6s)^sq205^*. *Tg(UAS:EGFP-2A-Δclocka-5×MYC)^sq215^*, abbreviated here as *Tg(UAS:EGFP-*Δ*clocka)* was generated by injection of *pT2-UAS:EGFP-2A-*Δ*clocka-5xMYC* using Tol2 mediated transgenesis. All experiments were carried out under guidelines approved by the IACUC of Biopolis (181408) and NTU (A19014). The minimal number of animals was used.

### 2.3 Two-photon Imaging

#### 2.3.1 Extended imaging

Based on the methods described by (Leung et al., 2019), non-anesthetized and non-paralyzed larvae were mounted in 2% low melting agarose (LMA), in a glass-bottom dish (Matek). The dish was filled with E3 medium supplemented with 1% Hepes and bubbled continuously with carboxygen (95% oxygen). Imaging was performed with a Nikon A1R MP system, attached to an upright FN1 microscope using a 25x 1.0 NA water dipping objective. Resonant scanning was performed with 2x averaging, and 20 focal planes were collected, spaced 5 μm apart. Signal was detected with a GaAsP detector. A piezo drive (Mad City Labs) was used for fast focusing during imaging. 21 stacks were collected, at an interval of 1 second, followed by a delay of 1 hour. The maximum imaging duration was 29 hours. After imaging, larvae were checked for blood flow. They were then released from the agarose. Only data from fish with normal blood flow and active swimming upon release were used for analysis.

To obtain a measure of calcium levels in the whole habenula, fluorescence intensities were summed at each time point using Fiji. The intensity at each hour was averaged and z-scores were plotted as a function of *zeitgeber* time.

#### 2.3.2 Short duration imaging

6-day-old *Tg(elavl3:H2B-GCaMP6s)* or *Et(−1.5hsp70l:Gal4-VP16)^s1011t^, Tg(UAS:GCaMP6f)* larvae were immobilized with mivacurium, mounted in 2% low melting temperature agarose and imaged with the two-photon system described above, using resonant scanning at a single plan. Frame size was 512×256 pixels and 2x averaging was used, leading to a frame rate of 14.5 Hz. Segmentation of cells expressing cytoplasmic GCaMP6f was performed manually in Fiji, while segmentation of cells expressing nuclear GCaMP6s was performed automatically with Suite2p (Stringer and Pachitariu, 2019). Z-scores were obtained for each cell in a recording, and cells were clustered using K-means clustering in Python. The classification was performed using maximum likelihood and leave-one-out cross-validation. Specifically, at each iteration we removed one dataset from the group and constructed empirical cluster distributions for both categories (ZT3 and ZT15) from the remaining datasets. The removed dataset was then assigned to the category whose empirical distribution maximized its likelihood. This process was repeated independently until all datasets were classified.

### 2.4 Immunohistochemistry and imaging of EGFP-ΔCLK

The *Tg(gng8:GAL4,UAS:EGFP-*Δ*clocka)* larvae at the age of 7 dpf were fixed in 4% PFA at 4°C for ~22 h. For immunostaining of extracted brains, skin covering the brain was manually removed and exposed brains were washed 3 times in PBST. Antigen retrieval was performed by incubating the fish in 150 mM Tris-HCl (pH 9.0) for 5 min at room temperature in order to equilibrate, and by subsequent heating of the samples at 70°C for 15 min. Following several washes and a blocking step, samples were incubated in the rabbit anti-GFP primary antibody (Torrey Pines Biolabs Inc., #TP401), diluted 1:500, and the mouse anti-Myc antibody (9E10; Santa Cruz Biotechnology), diluted 1:100, at 4°C for a few days. The secondary antibodies used were Alexa Fluor 488 goat anti-rabbit (ThermoFisher and Abcam), diluted 1:1000, and Alexa Fluor 647 goat anti-mouse (ThermoFisher), diluted 1:500. The immunostained brains were mounted in 2% LMA in PBS and imaged using a Zeiss LSM800 confocal microscope equipped with 40X 0.7 NA water immersion objective.

### 2.5 Screening and selection of *Tg(gng8:GAL4,UAS:EGFP-Δclocka)* fish

All of the *Tg(gng8:GAL4,UAS:EGFP-*Δ*clocka)* fish used in the following experiments were first screened for GFP fluorescence at 3 dpf using a Olympus MVX10 stereo fluorescence microscope. Since the expression of EGFP-ΔCLK in the habenula was mosaic, only individuals with expression in the majority of habenula cells were selected and used for further experiments as ΔCLK-positive fish. Individuals with zero expression as well as larvae exhibiting a minimal number of randomly labelled cells (less than ~15 cells per habenula) were used as the control siblings (Figure S1). Following the behavioural experiments, larvae were immobilized in tricaine methanesulfonate (MS-222; final concentration of 0.01 mg/ml; Sigma), mounted in 2% LMA in E3 medium and the habenula-specific expression of EGFP was assessed using a Zeiss LSM800 confocal microscope and 40X water dipping objective.

### 2.6 Quantification of DA and 5-HT levels

Three-month old female ΔCLK fish and their control siblings were immobilized using iced water and euthanized by decapitation at ZT3 and ZT15. Whole brains, excluding the olfactory bulb, were extracted in the ice-cold Ringer’s solution (116 mM NaCl, 2.9 mM KCl, 1.8 mM CaCl2, 5.0 mM HEPES, pH 7.2), snap-frozen in liquid nitrogen and stored in −80°C until further analysis. Supernatant was collected as described previously (Chatterjee and Gerlai, 2009). Quantification of dopamine (DA) and serotonin (5-HT) was carried out by LC/MS using the following analytical standards: dopamine hydrochloride (D2960000: Sigma-Aldrich), dopamine-1,1,2,2-d4hydrochloride (73483: Sigma-Aldrich) and serotonin (14927: Sigma-Aldrich). Three biological replicates (i.e. measurements of different samples) were performed.

### 2.7 Behaviour assays

#### 2.7.1 Acoustic startle following exposure to Schreckstoff

Thirteen to sixteen day-old ΔCLK larvae and their non-expressing siblings were placed in two transparent plastic tanks with dimensions of 80 × 54 × 33 (L × W × H in mm) and their locomotor activity was recorded from above at 10 fps using a camera (acA2040-90μm USB 3.0; Basler). Each of the two tanks was filled with 20 mL of facility water and held 8 fish at most. The water level in each tank was 5 mm in height to minimize vertical movement. The alarm substance was freshly prepared from adult zebrafish each week (Mathuru et al., 2012), and was tested for effectiveness on adults before use on larvae. An Arduino Grove Vibration Motor was positioned in between the two plastic tanks, on the stage holding the tanks, to provide the stimulus. The vibration was 1 sec in duration and was presented 3 times, at 1 minute intervals, starting five minutes before and after the delivery of the *Schreckstoff*. A startle response was defined as a change in body orientation of more than 90 degrees within 5 sec in response to each vibration tone. The response (yes or no for each fish) was averaged across 3 tones for each group to obtain a response percentage before and after the delivery of the *Schreckstoff*. Both the vibration tones and the delivery of *Schreckstoff* were controlled by an Arduino board synched with the video-recording Python codes. Experiments were performed on 3 separate batches of larvae. A different batch of freshly prepared skin extract was used for each experiment; the extract was verified for effectiveness by testing on adult fish. Only extract that evoked freezing was used.

#### 2.7.2 Locomotor activity

The *Tg(gng8:GAL4,UAS:EGFP-*Δ*clocka)* larvae were raised under standard conditions. Each experimental run involved animals from the same batch, originating from one pair of parents. The larvae were screened and selected as described in the previous section. At the age of 6 dpf the larvae were placed individually in a 24-well plate filled with 1.5 ml of tank water per well. The plate was then transferred into an incubator equipped with an IR camera (acA2040-90μm USB 3.0: Basler), a white LED box and four IR LED bars (LBS2-00-080-3-IR850-24V, 850 nm: TMS Lite). The camera and the LEDs were connected to a PC via a microcontroller board (Arduino UNO SMD). At the onset of recording (ZT6 – 9), fish were kept under ambient light, which was switched off at ZT14 and was kept off during the next day for the monitoring of activity under DD. A customized script written in Python 2.7, incorporating functions from the OpenCV library was used to control the Arduino microcontroller and simultaneously video-track fish locomotor activity. The video was captured using 576 × 864-pixel resolution at 10 frames per second and the background subtraction method was implemented to extract the ‘x’ and ‘y’ coordinates of each fish online. The position data was then converted to total distance (mm) travelled by each fish per 1 min time. The data conversion, further analysis and plotting were carried out using custom-written Excel macros, Python 2.7 and Estimation Statistics (Ho et al., 2019).

#### 2.7.3 Arousal threshold

Six days-old ΔCLK larvae and their control siblings were randomly placed in a 24-well plate and their locomotor activity was video-tracked as described in the previous section. In order to assess the arousal during sleep, vibration stimuli were applied through a speaker controlled, via an Arduino microcontroller board, by a modified version of the activity-tracking code in Python 2.7. Starting at ZT17, eighteen stimuli of nine different intensities, where 50% of computer audio output represented the highest intensity, were delivered for 200 ms (starting from 0%) in 12-16% increments every minute, first in a descending and then ascending order. The time interval between stimuli was determined based on previous findings that suggest a 30-second inter-stimulus gap is sufficient to avoid behavioural habituation (Burgess and Granato, 2007). The protocol was repeated at the commencement of every hour until ZT22, while the onset by ascending or descending order was altered each time. In total, 12 replicates (i.e. technical replicates) of each stimulus intensity were generated. A fish was scored as responding if it moved more than 7 pixels (~ 4 mm) following the stimulus delivery and the average percentage of responding fish was calculated for each of the stimulus intensities (Woods et al., 2014). The observed values were corrected for variations in the baseline locomotor activity, by first calculating the background probability of locomotion as a percentage of larvae which moved more than 7 pixels 5 s before the application of each stimulus. The average of 108 measures of the background probability of locomotion was then used to calculate the corrected probability of response at each of the stimulus intensities, implementing the following equation: corrected probability of response = observed value x [(observed value - background offset) / (max observed value - background offset)].

#### 2.7.4 Novel tank assay

All experiments were conducted between ZT2 and ZT4 on 3-5 month-old adult zebrafish, as described elsewhere (Haghani et al., 2019). Fish were recorded in pairs in a darkened room with a black background, where tanks were illuminated from the top with a natural light LED bar. Novel tanks were made of glass measuring 20 cm (L) by 5 cm (W) by 12 cm (H) with opaque sides (to prevent fish from viewing each other), placed side by side. Fresh water from the facility was added to tanks up to the 10 mL mark to make a total volume of 1 L. The setup was placed within a curtained area to obscure the experimenter.

Prior to exposure to the novel tank, fish were handled carefully to ensure that handling stress was minimized. Fish were gently netted in pairs from their home tanks in the facility to smaller crossing tanks, where they were brought into the room with the behavioural setup. Then, fish were netted into 100 mL beakers with ~30 mL of fish facility water and quickly poured into the novel tank. The recording was started within 30 s of fish being in the recording tanks. For exposure to *Schreckstoff*, fish were placed in 100 mL beakers containing 200 μL of heated skin extract (Chia et al., 2019) in 50 mL fish facility water for 2 minutes. Fish were netted into another beaker containing fresh facility water before being poured into the novel tank. Videos were acquired at 10 frames per second for 600 s (10 mins) using a Basler Ace camera (acA1300–200 μm; 1,280 × 1,024). The position (x-y coordinates) of the fish were tracked, and speed was binned by mm/s into a probability density curve.

For acute exposure to *Schreckstoff*, fish were first habituated in the glass tanks for 5-7 mins. Then, fish were recorded in the tanks for 5 mins before 1 mL of 1:4 dilution of heated skin extract was pipetted into the tanks. The recording continued for 5 mins after *Schreckstoff* delivery for the alarm response to be observed. Two fish were tested at one time, each in a separate tank, with one tank containing ΔCLK fish and the other a non-expressing sibling. The tank used for a particular genotype was switched at every trial. Characteristic alarm behaviours were quantified, such as time spent in the bottom third of the tank, number of darting episodes (fish speed >5 SD above average swim speed for the first 5 mins), and number of freezing episodes (movement of <5.5 mm per second). Tracking was automated to prevent observer bias. Sample size was based on previous experiments (Chia et al., 2019).

#### 2.7.5 Light/dark choice assay

All experiments were performed between ZT2 and ZT9 in a behavioural arena within blackout curtains as described previously (Cheng et al., 2016). Four transparent plastic tanks (dimensions: 43 mm W × 60 mm L × 30 mm H) filled with 30 mL fresh tank water served as assay chambers. Opaque cardboard sheets were placed between the 4 tanks. Tanks were placed on an Apple iPad screen and videos (9 fps) were recorded on a USB3.0 Basler camera (Model# acA2040-90umNIR 1440 × 1080 pixels) with a long-pass filter (MIDOPT, LP830 830 nm) placed above. Two IR light bars (850 nm peak from TMS-lite) positioned next to the 4 tanks served as an IR light source. The light/dark compartments were created by white and black rectangles (with 50% transparency level for the black) on a Microsoft PowerPoint slide displayed on the iPad Air at highest brightness level. Sample size was based on previous experiments(Cheng et al., 2016).

Larvae were exposed either to *Schreckstoff* or to control water (embryo medium) in a treatment tank of the same dimensions with the same volume of tank water for 10 minutes. Larvae were then gently pipetted from the treatment tank into two wash-out tanks sequentially before being placed into the dark/light test chamber. Real-time tracking of the fish was achieved by custom-written Python codes using the OpenCV library and recorded as coordinates of the centroid of the fish across time. Larval behaviour was recorded for 10 minutes. Offline analysis was performed using custom-written Excel macros to determine the position of the animals, the number of entries into each compartment, the percentage of total time spent in each compartment, etc. A few fish (N=2 in both the Control group and Gng8/ΔCLK group) were excluded from the analysis because they were entirely immobile or because of tracking errors.

### 2.8 In situ hybridization

Hybridisation chain reaction with split initiator probes was conducted using the HCR v3.0 protocol for whole-mount zebrafish embryos and larvae from Molecular Instruments. Probe hybridization buffer, amplification buffer, probe wash buffer, probe sets and HCR amplifiers were obtained from Molecular Instruments.

*Tg(gng8:GAL4, UAS:EGFP-*Δ*clocka)* larvae were maintained at 28°C under the standard 14-10h LD cycle. For characterization of clock genes under DD, nacre and *Tg(gng8:GAL4, UAS:EGFP-*Δ*clocka)* larvae were maintained under standard LD conditions transferred to a dark room at ZT14 the day before fixation.

All *Tg(gng8:GAL4, UAS:EGFP-*Δ*clocka)* larvae were screened for mosaic expression of EGFP-ΔCLK at 5 dpf and EGFP-ΔCLK positive larvae were separated from non-expressing siblings. All larvae were fixed at ZT3, ZT7, ZT11, ZT15, ZT19 and ZT23h with 4% paraformaldehyde (PFA) in phosphate buffered saline (PBS) at 4°C in a 1.5mL eppendorf tube (1 tube per time-point) and incubated at 4°C overnight. Fixations for DD larvae were performed under dim light. For dehydration and permeabilization, larvae were washed 4 × 10 minutes and 1 × 50 minutes with 1mL of 100% methanol (MeOH) at room temperature, and subsequently stored in −20°C for several days. For rehydration, larvae were washed 1 × 5 minutes with 1mL of 75% MeOH (in PBST), followed by 50%, and 25% MeOH (in PBST), and subsequently 5 × 5 minutes washes with 100% PBST. Larvae were then treated with 1mL of 30μg/mL proteinase K for 45 minutes at room temperature and washed twice with 1mL of PBST without incubation. Postfixation of larvae was conducted by adding 1mL of 4% PFA for 20 minutes at room temperature. The larvae were then washed 1 × 1 hour and 4 × 5 minutes with 1mL of PBST. For pre-hybridization, 4 larvae from each sample were transferred to a 1.5 mL eppendorf tube and 500 μL of probe hybridization buffer was added and incubated at 37°C for 30 minutes. For detection of clock genes, probe hybridisation buffer was aspirated from the samples and probe solution (2 μL each of 1 μM *arntl1b* and *per3* qHCR probe sets or 3 μL of *cry1a* dHCR probe set in 500 μL of probe hybridisation buffer at 37°C) was added and incubated at 37°C for ~22 hours. Samples were washed 4 × 15 minutes with 500 μL of probe wash buffer at 37°C, followed by 2 × 5 minutes washes with 500 μL of 5X SSCT at room temperature for removal of excess probes. 5X SSCT was aspirated from the samples and 500 μL of amplification buffer was added and incubated for 30 minutes at room temperature. For amplification of signal, amplification buffer was aspirated from the samples and hairpin solution (10 μL each of snap-cooled HCR amplifier hairpins h1 and h2 for all 4 probe sets in 500 μL of amplification buffer at room temperature) was added and incubated in a dark box at room temperature for ~22 hours. Hairpin solution was then removed by 2 × 5 minutes and 2 × 30 minutes washes and another 1 × 5 minutes wash with 500μL of 5X SSCT at room temperature. Samples were lastly stored at 4°C in a dark box before microscopy.

#### 2.8.1 qHCR Imaging

4 larvae from each sample were mounted in 2% LMA in PBS in dorsal view and imaged using a Zeiss LSM800 upright confocal microscope with 40 × water dipping objective at 0.5x zoom. The acquisition settings were identical in all experiments. Z-stacks were collected at 512×512 resolution, giving a x-y pixel size of 0.62 μm, and a z-step of 0.73 μm. Sample size was based on (Trivedi et al., 2018).

#### 2.8.2 Image Analysis

To quantify gene expression in the entire habenula across different fish, segmentation of habenula was performed manually using the Segmentation Editor in Fiji (Schindelin et al., 2012). The integrated intensity of *per3* and *arntl1b* was measured using the 3D Objects Counter (Bolte and Cordelières, 2006). The ratio was calculated to compensate for potential tube-tube variation as well as change in intensity with imaging depth. Values were plotted using GraphPad Prism.

To compare the level of *per3* in ΔCLK expressing and non-expressing cells, within-fish analysis was performed on single planes. ΔCLK cells were segmented by thresholding the GFP signal, while the habenula was manually segmented. Masks for control cells were obtained by subtracting the ΔCLK mask from the habenula mask. ROIs were then created, and mean intensity measured, based on methods described by (Choi et al., 2018), i.e. with a mean filter of 3 and background correction. Statistical analysis was performed with estimation statistics (Ho et al., 2019). 5000 reshuffles of the control and test labels were performed to calculate each permutation P value.

#### 2.8.3 dHCR imaging

Following label for *cry1a*, 3 larvae from each of 6 time-points were imaged with confocal microscopy, using an Achroplan 63x water dipping objective and 2048 × 4093 image size, giving a x-y pixel size of 0.0495 μm. z step was 0.78 μm. Sample size was based on Choi et al, 2018. To analyse the images, the habenula in one plane ~ 10 μm from the surface was manually segmented. Spots were identified in Fiji using a Laplacian transform with FeatureJ, with the smoothing scale set at 5. The number of spots was counted using FindMaxima, with the prominence > 10.0. Identical settings were used for acquisition and analysis of all dHCR images.

## 3. Results

### 3.1 Multiple clock genes are expressed in the zebrafish habenula

Using in situ hybridization in brain sections with DIG-labelled probes, a number of clock genes, such as *per1b* and *bmal* (*arntl1a*), have been shown to be expressed in the habenula of adult zebrafish (Weger et al., 2013). To better characterize clock gene expression in this structure, we analyzed single cell transcriptomes of the zebrafish habenula (Pandey et al., 2018). A diversity of clock gene components, including *arntl1b*, *arntl2*, *per1a*, *per2*, *per3*, *cry1a*, *cry1ab*, *cry1ba*, *cry1bb*, *clocka*, *clockb* and *nr1d1* were found (Figure 1). These transcripts were isolated from *gng8*-expressing habenula neurons, which are found throughout the habenula (Hong et al., 2013; Pandey et al., 2018). All clusters contained clock genes, suggesting that the molecular clock is broadly distributed.

**Figure 1.**
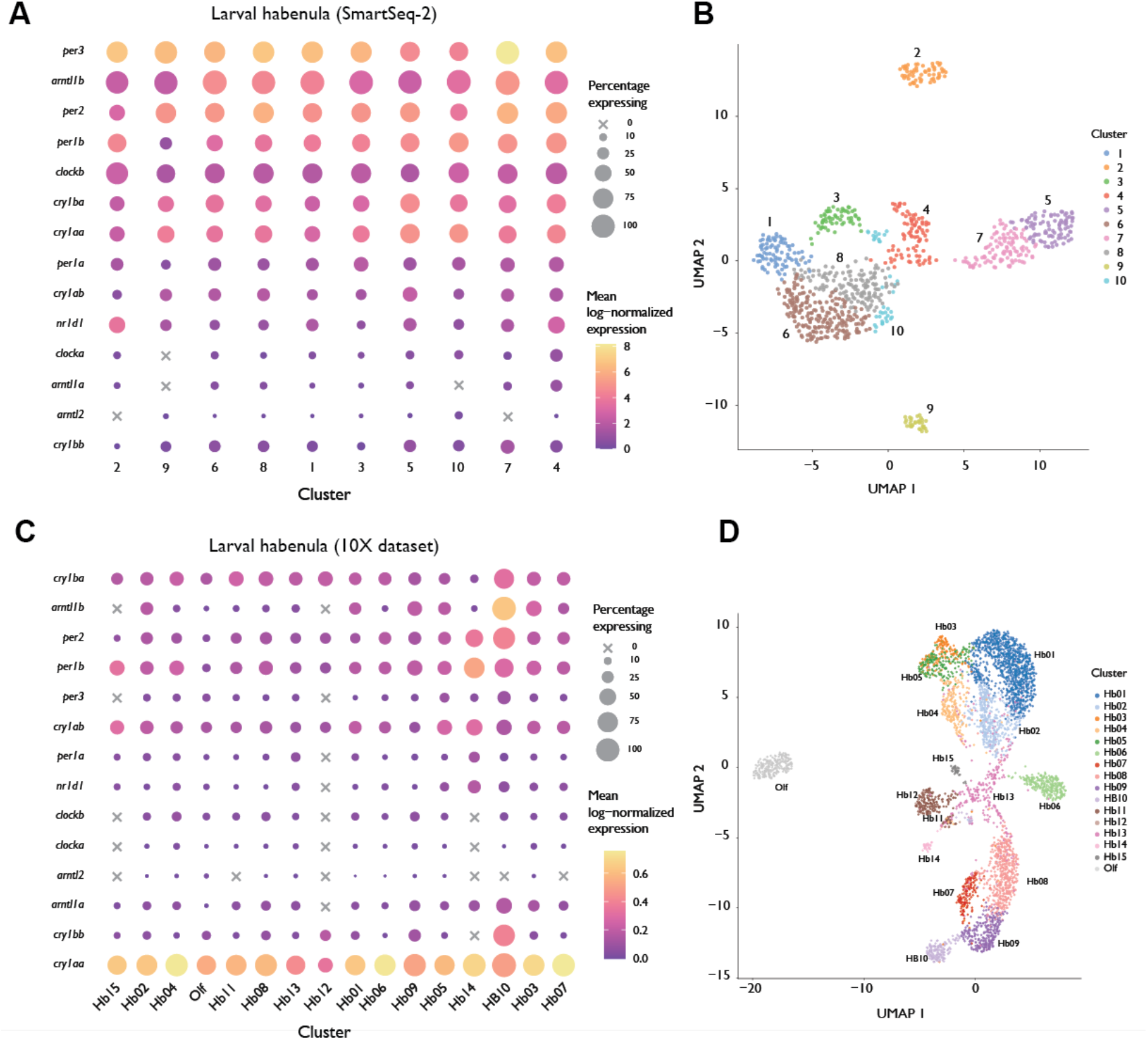
Clock genes expression in the zebrafish habenula based on single cell RNA sequencing. **A)** Dot plot showing mean log-normalized expression per cluster and percentage of cells per cluster expressing each gene in the SmartSeq-2 dataset from Pandey et al. **B)** t-SNE representation of clustering results in the SmartSeq-2 dataset. Each point is a cell. **C)** same as A) but in the 10X dataset. **D)** same as C but in the 10X dataset. The cluster assignment for the 10X dataset corresponds to those assigned by Pandey et al 2018.

To examine the temporal dynamics of the clock in larval zebrafish, quantitative in situ hybridization using hybridization chain reaction (HCR) (Choi et al., 2018) was carried out on fish that were fixed at 6 different time points (Figure 2). With qHCR imaging, *arntl1b* appeared to be expressed at a higher level in the habenula compared to the rest of the brain (Figure 2A), with a stronger signal at approximately *zeitgeber* time ZT 15; expression at ZT 19 and ZT 23 was detected at the posterior margins. *per3* was detectable broadly in the brain, including the entire habenula, with strongest expression at ZT 23 and ZT 3 (Figure 2B). Signal intensity obtained with qHCR imaging provides a relative measure of transcript level (Choi et al., 2018), and can vary with sample thickness and between tubes. To minimize error from these potential confounds, we calculated the ratio of *arntl1b/per3* as a means of comparing gene expression between animals. This ratio showed a circadian variation, with a peak at ZT 11 - 15 (Figure 2D). To test whether clock gene cycling requires light, we examined expression in fish grown in constant darkness. The ratio of *arntl1b*/*per3* expression was maximal at CT 15 and minimal at CT3 (Figure 2E-G). We used single molecule imaging, with dHCR imaging of *cry1a*, to further test whether the clock is cycling. Again, a circadian change in expression level was detected in the habenula in constant darkness (Fig 2H, I), with a peak at CT3 and a trough at CT15, similar to what has been reported in other tissues under LD (Cavallari et al., 2011) or DD conditions (Kobayashi et al., 2000). These observations confirm previous findings using clock reporters (Wang et al., 2020; Weger et al., 2013) that the molecular clock is active in the zebrafish habenula.

**Figure 2.**
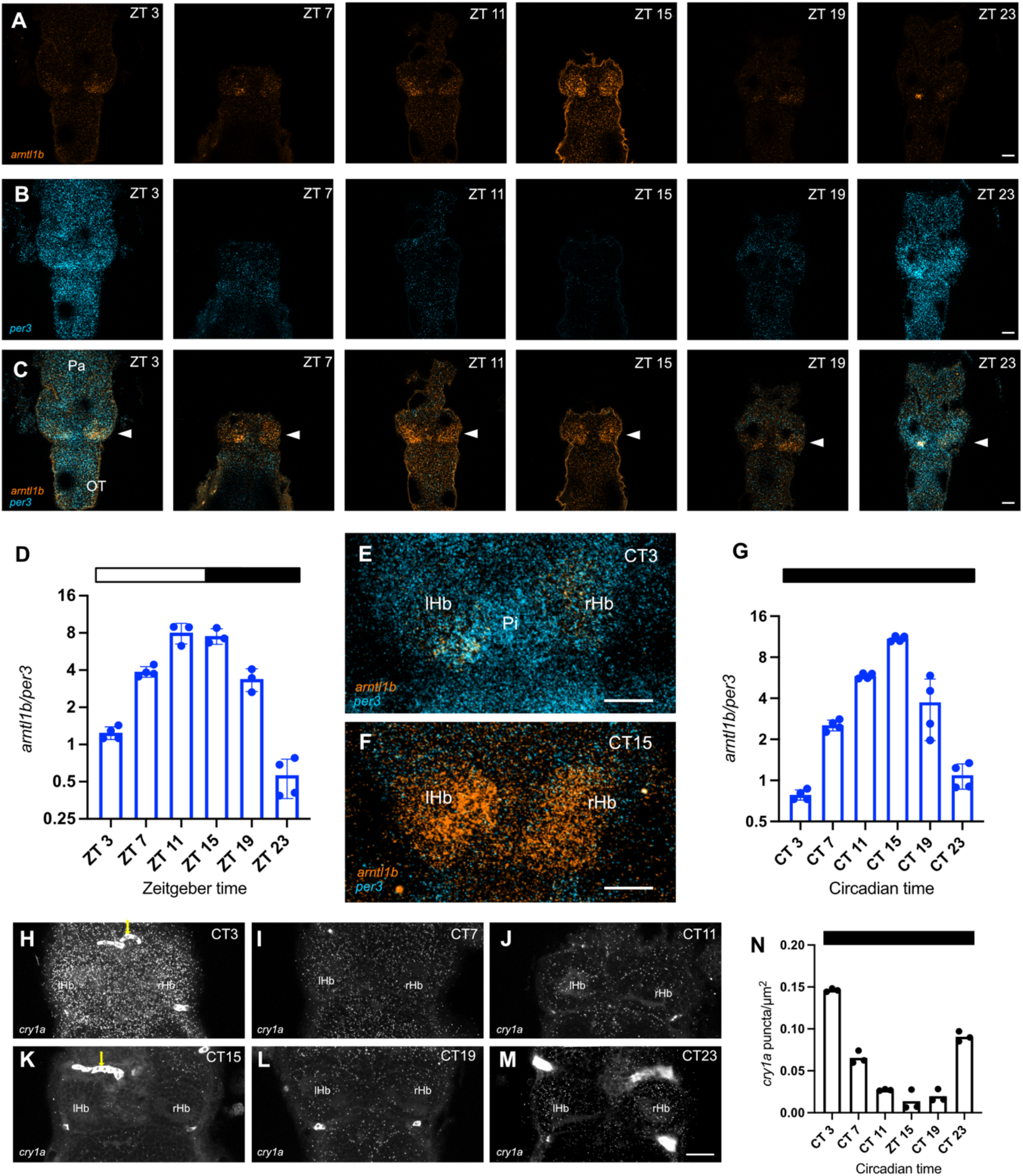
Dynamics of clock gene expression in the zebrafish habenula. **(A-C).** Single optical sections of the brain of 6 day old larvae fixed at six different time points, labelled for *arltn1b* (A) and *per3* (A) using hybridization chain reaction. **(C)** Merge of (A) and (B). Arrowheads indicate the habenula. (**D**) Ratio of *arntl1b/per3* in the habenula of fish in a normal LD cycle. (**E-G**) Expression of *per3* and *arntl1b* in the habenula of fish grown under DD conditions. (**H-M**) *cry1a* expression in the habenula of fish grown under DD conditions. Images in E and F are maximum projections of the entire habenula. Images in panels H-M are maximum projections spanning 5 μm. The arrowheads indicate autofluorescent blood cells. A, B, C, E, F, H-M are dorsal views with anterior to the top. The dots in panels D, G and N indicate values for individual fish. The horizontal bars indicate illumination conditions. lHb: left habenula; rHb: right habenula; Pi: pineal; Pa: pallium; OT: optic tectum. Scale bar = 25 μm.

### 3.2 The zebrafish habenula displays circadian variation in intracellular calcium

The presence of the molecular clock should lead to a number of circadian changes in a cell. One such change is the level of cytoplasmic calcium (Pennartz et al., 2002). To examine intracellular calcium levels in the zebrafish habenula, two-photon imaging was carried out in fish held in constant darkness. Confocal microscopy was not used to avoid light-induced disruption of the molecular clock by visible light (Tamai et al., 2007). By imaging a series of z-stacks every hour, such that the entire habenula could be captured (Figure 3A-D), we observed an increase in overall fluorescence intensity during the subjective day, followed by a decrease during the subjective night (Figure 3E; 5 out of 6 fish imaged). This finding is consistent with the hypothesis that the habenula contains an active circadian clock that has the potential to drive changes in the intracellular calcium even in the absence of light, a prominent circadian *zeitgeber*.

**Figure 3.**
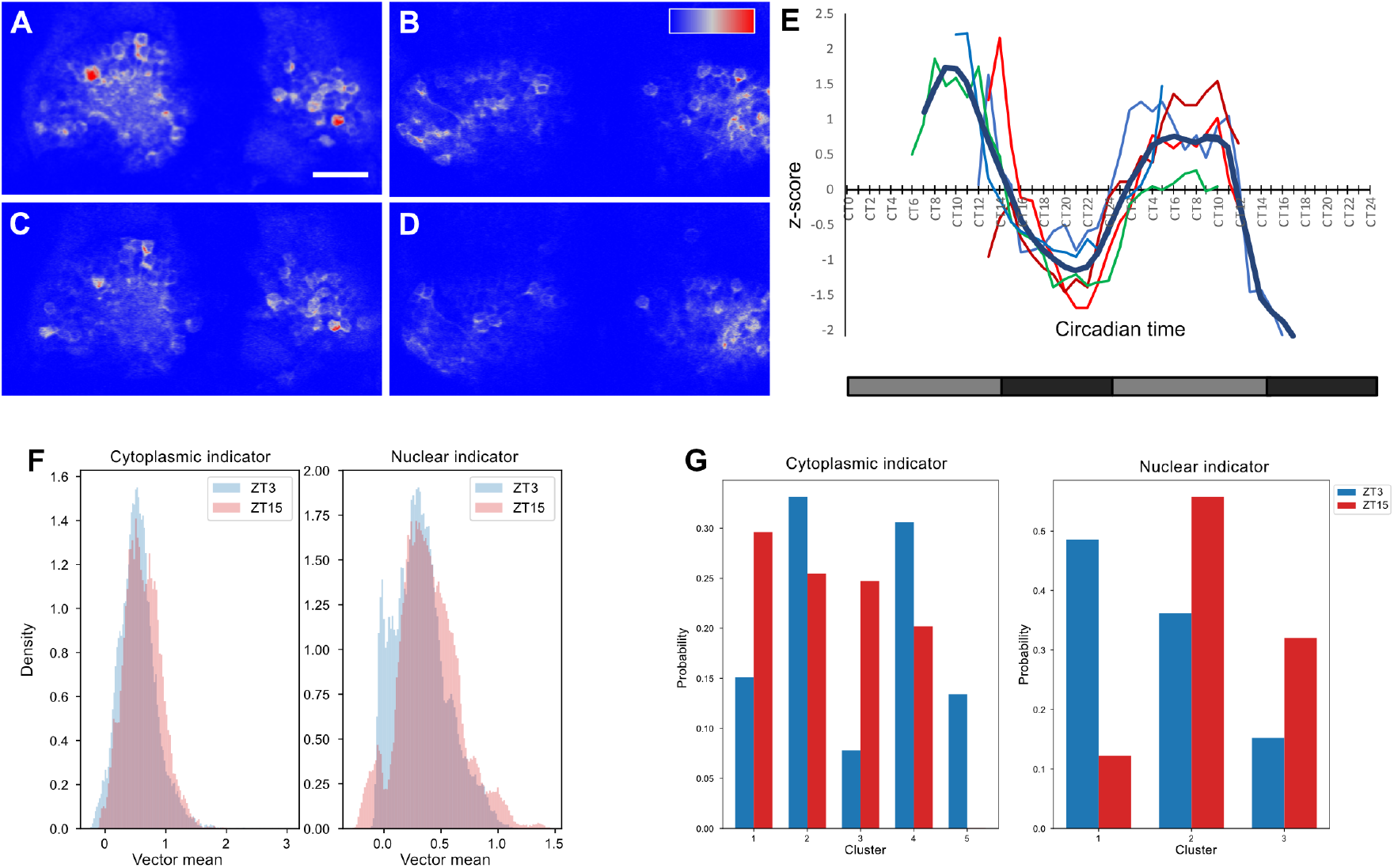
Circadian variation in cytoplasmic calcium in habenula neurons. (**A-E**) Long duration multiplane recording of the habenula of *Et(GAL4s1011t),Tg(UAS:GCaMP6s)* larvae in constant darkness. (A-D) Examples of imaging data from one fish at two different time points, ZT5 (A, B) and ZT14 (C, D), and two different focal planes, dorsal (A, C) and ventral (B, D). Scale bar = 25 μm, dorsal view with anterior to the top. (E) Average level of cytoplasmic calcium in the habenula, as shown by intensity of GCaMP6s. Each coloured trace represents one fish. The thicker black line is the mean. There is a reduction in intracellular calcium levels during the subjective night. (F-G) Difference in relative activity levels of habenula neurons between day (ZT3) and night (ZT15) as measured by both cytoplasmic and nuclear GCaMP6s. (F) Distribution of the mean of vectors containing z-scores of neurons with high variance in their activity. (G) K-means cluster distribution for the same vectors.

In the mouse, neurons in both medial (Sakhi et al., 2014a) and lateral (Sakhi et al., 2014b) habenula display spontaneous neural activity that varies across a circadian cycle. To assess whether this is also the case in the zebrafish habenula, we carried out calcium imaging at ~15 Hz at a single plane. We imaged at two different time points, starting at ZT3 and ZT15. The z-score for fluorescence of each cell was calculated, and a vector was generated for the activity of cells with high variance. A difference was observed between the means of this vector at ZT3 and ZT15 (Figure 3F). To further characterize the activity, cells were classified using K-means clustering. The cluster composition was distinct between day and night (Figure 3G). To test the robustness of this apparent difference, we asked if a classifier could correctly assign a random dataset to either ZT3 or ZT15. Indeed, 21 out of 26 datasets of the cytoplasmic indicator and 10 out of 13 datasets with the nuclear indicator were correctly assigned. These findings suggest that neural activity in zebrafish habenula is different at day versus night.

### 3.3 Expression of a truncated *clock* gene in the habenula affects function

The expression of clock genes and the circadian change in intracellular calcium in the habenula raises the possibility that an intrinsic clock influences habenula function. To test this, we selectively expressed a truncated *clocka* gene (Dekens and Whitmore, 2008), referred to as Δ*clocka*, in the habenula. This construct has been shown to act in a dominant manner to inhibit the molecular clock in the zebrafish pineal (Livne et al., 2016). Δ*clocka* was expressed in the habenula using the GAL4/UAS system with *TgBAC(gng8:GAL4)* (Hong et al., 2013) as the driver. In addition to Δ*clocka*, the effector construct also included enhanced green fluorescent protein (EGFP), which was separated from ΔClocka (ΔCLK) by a 2A peptide linker, thus enabling the expression of two separate proteins. The presence of EGFP in the habenula neurons of *Tg(gng8:GAL4, UAS:EGFP-*Δ*clocka)* fish (Figure 4A, B) indicates that the construct is expressed appropriately and enables convenient identification of ΔCLK-positive individuals (Livne et al., 2016). Labelling with an antibody against the Myc tag, which is fused in frame at the C-terminus of ΔCLK, indicates that the truncated protein is translated (Fig. 4C). Cells expressing ΔCLK would be expected to have a lower level of Clock-regulated genes such as *per*, than non-expressing cells. To assess this, qHCR imaging was performed on larvae with scattered expression of ΔCLK, and the level of *per3* in cells with EGFP expression was compared with the ratio in non-expressing cells in the same sample. As seen in Fig. 4D the level of *per3* differed between ΔCLK and control cells at all time points, with a paired Cohen’s D of 1.58 [95.0%CI 1.03, 4.22] at CT3, 0.525 [95.0%CI 0.337, 4.25] at CT7, −1.89 [95.0%CI −4.78, −0.734] at CT11, −1.57 [95.0%CI −3.38, −1.15] at CT15, 0.872 [95.0%CI 0.636, 2.26] at CT19 and 0.637 [95.0%CI 0.536, 0.816] at CT23. This suggests that expression of ΔCLK influences the molecular clock, although it does not eliminate cycling.

**Figure 4.**
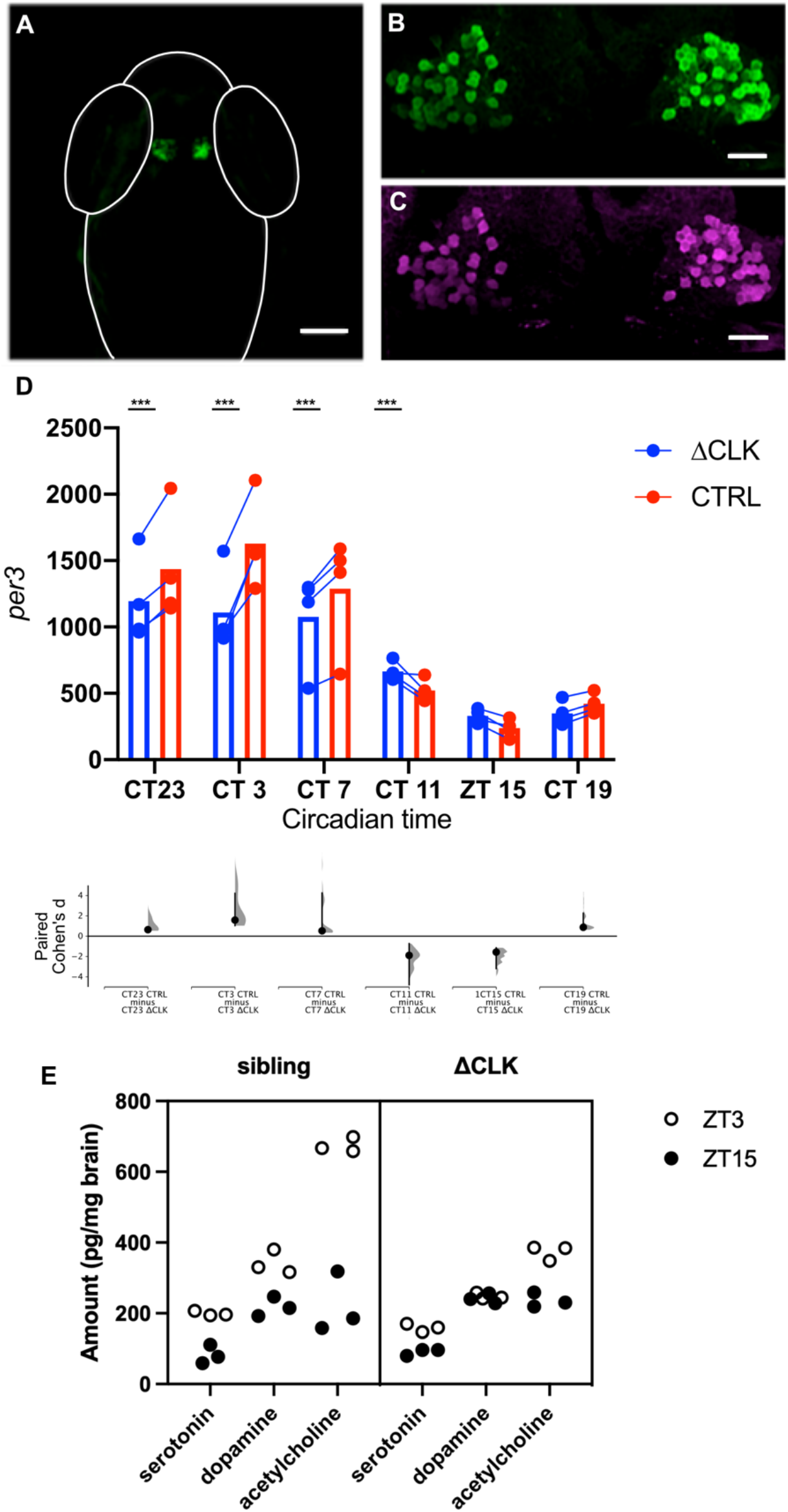
The effects of expressing truncated *clocka* in the habenula. (**A**) Specific expression of EGFP in the habenula of a *Tg(gng8:GAL4, UAS:EGFP-*Δ*clocka)* fish. (**B)** At higher magnification, EGFP is visible in the cytoplasm of habenula neurons. (**C**) Detection of Myc, which is fused to the C-terminus of ΔCLK, in the habenula of transgenic fish. **(D)** Level of *per3* in the habenula of fish with mosaic expression of EGFP-ΔCLK, as determined by qHCR imaging. The lower graph shows Cohen’s d, indicating the degree of difference between ΔCLK-expressing and non-expressing cells at each time point. Values are given in the main text. **(E)** Effects of ΔCLK expression in the habenula on global neuromodulator levels. Plot showing the amount of secreted serotonin, dopamine and acetylcholine in the whole brain of and *Tg(gng8:GAL4, UAS:EGFP-*Δ*clocka*) and sibling fish (N=3) at ZT3 and ZT15. Fish with ΔCLK expression in the habenula have a reduced day-night change in levels. Scale bar = 100 μm in panel A, 20 μm in panels B and C; *** indicates p<0.0002 by permutation t-test.

Given the partial inhibition shown by gene expression analysis, we asked whether is any evidence that expression of ΔCLK affects habenula function. The habenula regulates the broad release of serotonin and dopamine (Amo et al., 2010; Satar et al., 2020) and contains cholinergic neurons (Hong et al., 2013). If expression of ΔCLK has any effect on habenula function, the global levels of secreted serotonin, acetylcholine and dopamine, which vary in a circadian fashion (Hut and Zee, 2011; Mendoza, 2017), should be affected. To test this, brains were isolated from adult zebrafish expressing EGFP-ΔCLK and from their non-expressing siblings, at ZT3 and ZT15. The supernatant from extracts were analysed by liquid chromatography and mass spectrometry (LC/MS) (Chatterjee and Gerlai, 2009). While non-expressing fish had a clear difference in the day- and night-time levels of dopamine (Figure 4E; mean difference = 124 pg/mg, p < 0.0002 by permutation t-test), there was almost no difference in fish expressing ΔCLK protein (mean difference = 6.3 pg/mg, p = 0.30 by permutation t-test). The mean difference in between day and night-time levels of serotonin and acetylcholine levels was also higher in control fish (117 pg/mg and 454 pg/mg respectively, p < 0.0002 by permutation t-test in both cases), compared to ΔCLK-positive fish (68.1 pg/mg and 137 pg/mg respectively, p < 0.0002 by permutation t-test). This change of neuromodulator levels is consistent with the notion that expression of ΔCLK in the habenula affects function.

### 3.4 Habenula expression of ΔCLK reduces anxiety-like behaviour induced by the alarm substance

We hypothesized that a stress response would be affected by the expression of ΔCLK in the habenula. One such response for fish is the alarm response, which is triggered by the pheromone *Schreckstoff*. In adult zebrafish, Schreckstoff causes an acute change in swimming behaviour (Jesuthasan and Mathuru, 2008; Speedie and Gerlai, 2008). Following transient exposure to *Schreckstoff*, fish display abnormal scototaxis (Maximino et al., 2014) and heightened vigilance. To determine whether manipulation of the habenula clock influenced the acute response, adult fish were placed individually in a test tank. After a period of acclimatization, the alarm substance was introduced into the tank (Figure 5A). Adult *Tg(gng8:GAL4, UAS:EGFP-*Δ*clocka)* zebrafish showed an increase in darting (Figure 5C, paired mean difference of 1.33 seconds [95.0% CI 0.5, 2.67], p = 0.0260 Wilcoxon rank sum test), as did their non-expressing siblings (paired mean difference of 1.25 seconds [95.0% CI 0.417, 3.33], p = 0.0256 Wilcoxon rank sum test). There was no difference in the movement to the tank base (Figure 5D; paired mean difference of 16.4 [95.0%CI 1.02, 41.2], p = 0.530 for ΔCLK fish and 12.6 [95.0%CI −0.997, 28.1], p = 0.182 for siblings).

**Figure 5.**
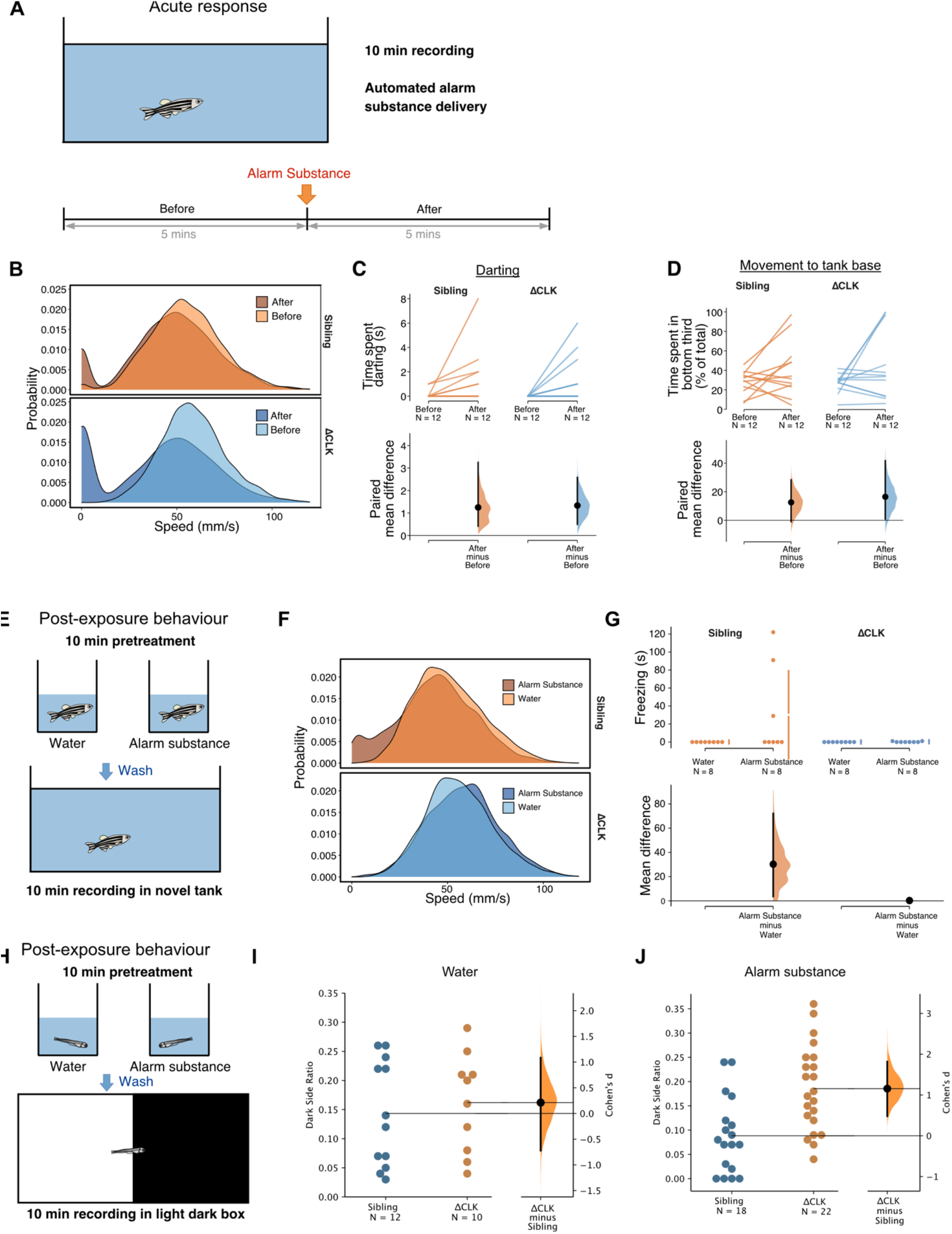
The effects of the alarm substance *Schreckstoff* on *Tg(gng8:GAL4, UAS:EGFP-*Δ*clocka)* fish. (A-D) Acute effects in adult fish (N=12). (A) Schematic of the experiment. (B) Speed distribution in fish before and after exposure to the alarm substance, where the substance remained in the tank after delivery. (C-D) Time spent darting (C), restricted to the tank base (D) in the presence of the alarm substance. As indicated by the paired mean difference plots (lower plot), fish with and without expression of truncated *clocka* behave similarly. (E-G) Post-exposure behaviour in adult fish (N=8). (E) Schematic diagram of the experiment. (F) Speed distribution in a novel tank containing clean water, after exposure to alarm substance or clean water. (G) Only sibling fish exhibited freezing after exposure to alarm substance. (H-J) Post-exposure behaviour in 2 week-old zebrafish. (H) Schematic diagram of the experiment. (I) Percentage of time in the dark area of a tank, after 10 minutes of exposure to embryo water. (J) Percentage of time spent in the dark side after exposure to the alarm substance. *Tg(gng8:GAL4, UAS:EGFP-*Δ*clocka)* fish spend more time in the dark side, with a Cohen’s D of 1.2.

To monitor expectation of danger (or anxiety-like behaviour), individual fish were exposed to the alarm substance in a beaker, then transferred to a novel tank containing fresh water (Figure 5E). ΔCLK fish, in contrast to their non-expressing siblings, did not freeze, but behaved similarly to fish that had been exposed to fish water (Figure 5F, G)). We also monitored the post-exposure response of larval zebrafish, using a light-dark choice assay as an indicator of stress (Steenbergen et al., 2011) (Figure 5H). Larval *Tg(gng8:GAL4, UAS:EGFP-*Δ*clocka)* fish displayed less dark avoidance after transient exposure to Schreckstoff, as indicated by the increased amount of time spent in the dark side of the tank (Figure 5I, J; mean difference of 0.0971 [95.0% CI 0.0475, 0.146], p = 0.001 permutation t-test; Cohen’s d = 1.16 [95% CI 0.481, 1.82]), in contrast to their non-expressing siblings (mean difference of 0.0213 [95.0% CI −0.058, 0.0928], p = 0.588; Cohen’s d = 0.224 [95% CI −0.667, 10.8]). Together, these observations suggest that disruption of the habenula clock leaves acute responses unaffected but reduces the expectation of danger that is evoked by transient exposure to the alarm substance.

We next asked whether the expectation of danger induced by the alarm substance varies in a circadian fashion in zebrafish; a circadian change has been noted in another species, *Rasbora heteromorpha* (Thines and Vandenbussche, 1966). To do this, we exposed larval fish to the alarm substance either at ZT3 or ZT15. Larval zebrafish respond to the alarm substance by extended immobility (Jesuthasan et al., 2020). To sensitively detect this, a vibration was presented five minutes before and after delivery of the alarm substance (Figure 6A). Control zebrafish showed a reduction in startle after delivery of the alarm substance during the day (ZT3, p = 0.00248, Pearson’s chi-squared test in R), but not at night (ZT15, Figure 6B), consistent with freezing during the day but not at night. *Tg(gng8:GAL4, UAS:EGFP-*Δ*clocka)* fish did not show a reduction in startle at day or at night (Figure 6C). These data suggest that the extended freezing evoked by the alarm response is circadian in zebrafish and is influenced by the habenula clock.

**Figure 6.**
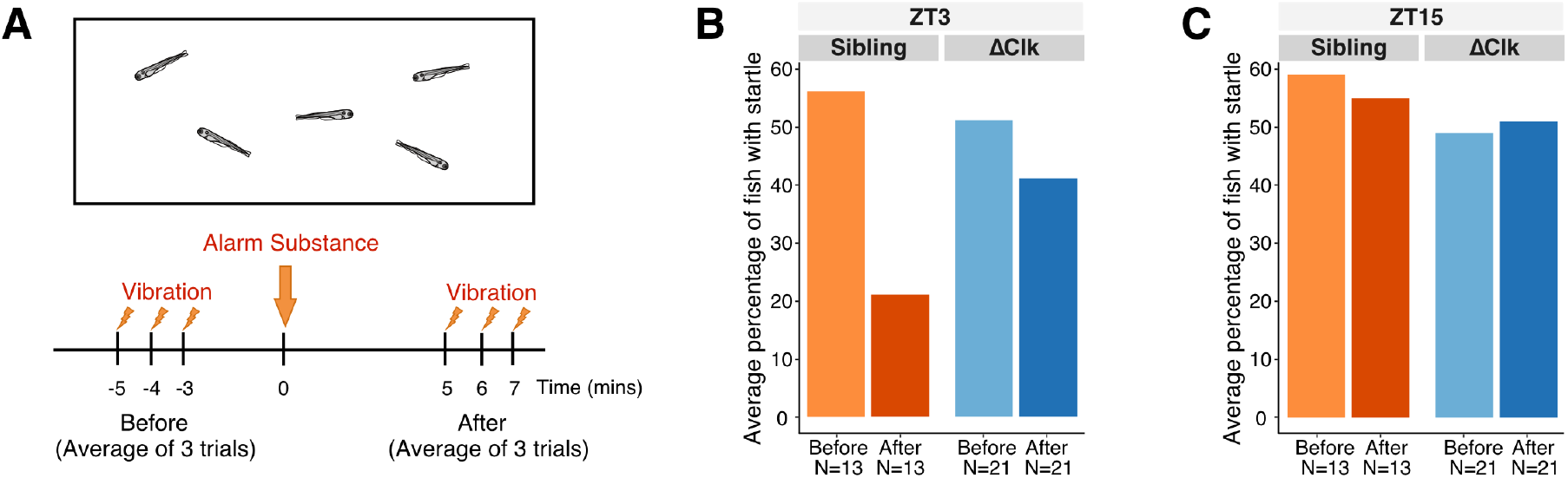
Day-night variation in the response of zebrafish to *Schreckstoff*. (A) Schematic of the experiment. To assess freezing in larval fish following exposure to the alarm substance, fish were exposed to vibrations in the day (ZT3) (B) or at night (ZT15) (C). Control fish, which are non-expressing siblings of *Tg(gng8:GAL4, UAS:*Δ*clocka)* fish, show a reduction in startle after exposure during the day, but not at night. This response to alarm substance was not seen in *Tg(gng8:GAL4, UAS:*Δ*clocka)* fish in the day or night.

### 3.5 Expression of ΔCLK in the habenula does not affect pineal-regulated circadian behaviour

We hypothesized that while the circadian clock in the habenula affects the response to a stimulus that influences expectation, circadian behaviours that are under the control of the pineal clock (Livne et al., 2016) should be unaffected. To test this, we analysed the rhythmic locomotor activity of ΔCLK-positive larvae and their non-expressing siblings. All larvae were entrained to the standard LD cycle and their locomotion was monitored for a few hours during the light phase, followed by the onset of darkness at ZT14 for activity tracking during the night. Subsequently, the fish were monitored for one day under constant darkness (DD), in order to eliminate the masking effects of light and dark on clock-regulated behaviour (Livne et al., 2016). As seen in Figure 7A-D, ΔCLK larvae showed a similar pattern of locomotion as their non-expressing siblings, becoming mobile during the light phase and during the subjective day, while exhibiting quiescent periods of inactivity during the 1st and 2nd night of the recording.

**Figure 7.**
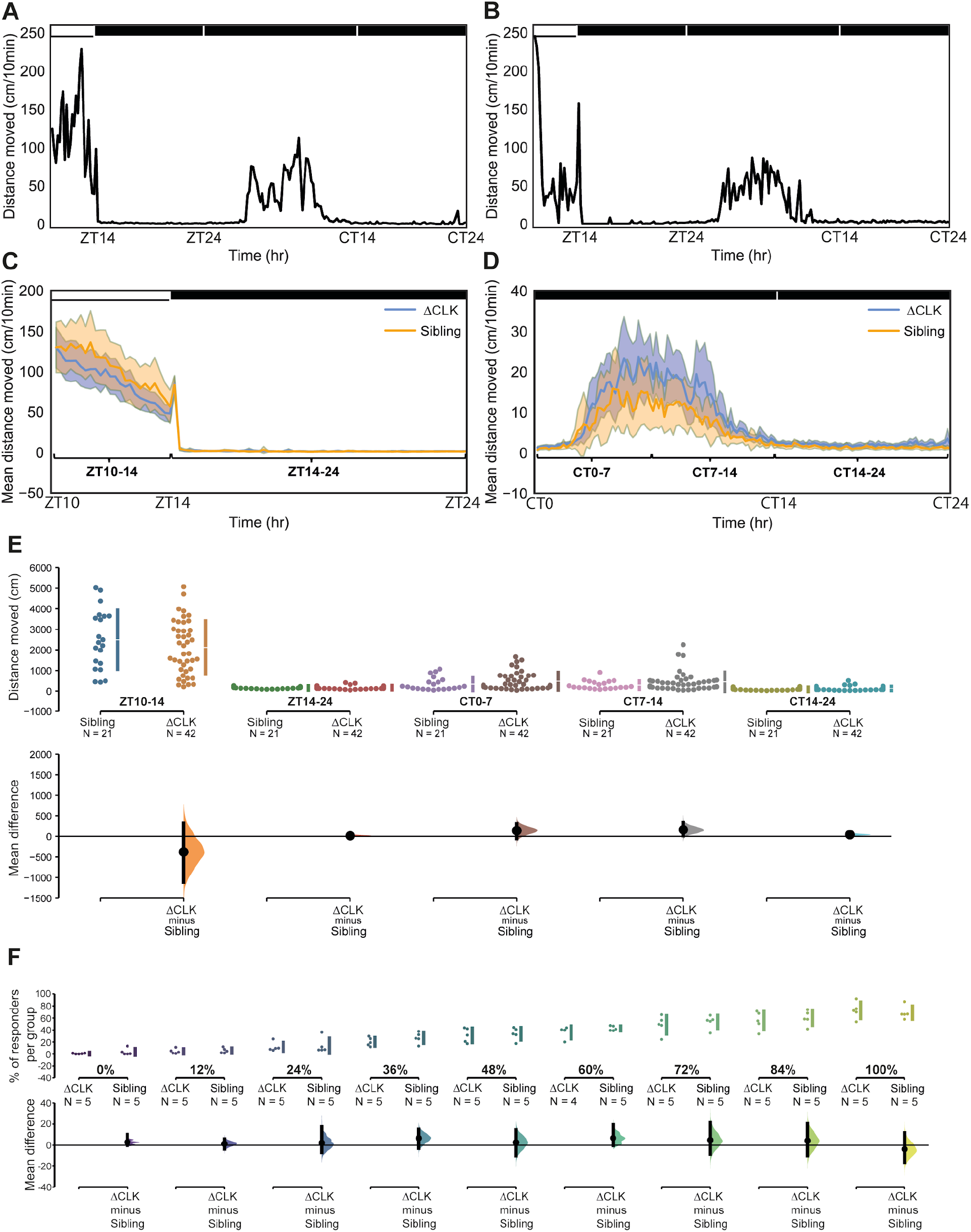
Sleep is unaltered in *Tg(gng8:GAL4, UAS:EGFP-*Δ*clocka)* larvae. (**A, B**) Distances moved (cm/10min) by individual larvae from (A) Sibling group and (B) ΔCLK group are plotted against time (hr) over the time-course of the whole recording. CT = circadian time. (**C, D**) Mean distances moved (cm/10min) of ΔCLK (blue) and Sibling (orange) groups are plotted against time (hr) (C) during the first 14 hours of recording, including 4 hours of light-phase and 10 hours in the dark during the night and (D) during the subsequent 24 hours of recording under DD. 95% CIs are indicated by the shaded areas; horizontal brackets indicate 5 analyzed timeframes. Black and white bars represent periods of dark and light, respectively. (**E**) The Cumming estimation plot shows the mean differences in 5 timeframes of the recording: ZT10-14 (light-phase), ZT14-24 (1st night), CT0-7 (early subjective day), CT7-14 (late subjective day) and CT14-24 (subjective night). The upper axis shows the raw data, with each dot representing the overall distance moved (cm) by a single larva; gapped vertical lines to the right of each group indicate mean ± SD. Each mean difference is plotted on the lower axes as a bootstrap sampling distribution. Mean differences are depicted as black dots and 95% CIs are indicated by the end of vertical error bars. (**F**) Analysis of sensory responsiveness of the *Tg(gng8:GAL4, UAS:EGFP-*Δ*clocka)* larvae (7 dpf) during the delivery of auditory stimuli of 9 different intensities. The Cumming estimation plot shows the mean differences for 9 comparisons, corresponding to 9 stimulus intensities (0-100%). The raw data is plotted on the upper axis, each dot representing the percentage of responders per single experimental run.

In addition to locomotion, we also examined arousal threshold, which is normally elevated during the sleep-like state at night. Arousal threshold was measured by responsiveness to auditory stimuli (Zhdanova et al., 2001). Nine different intensities were delivered between ZT17 and ZT23, and a sudden change in locomotion was taken as an indicator of responsiveness. A larva was scored as a responder if it moved more than 7 pixels after the stimulus and the average percentage of responders was calculated for each stimulus intensity per group for each experimental run (N = 5 experimental runs). Overall, no difference was seen between ΔCLK larvae and their control siblings (Figure 7E, F). Together, these experiments indicate that disruption of the habenula clock has relatively specific effects.

## 4. Discussion

Predictive coding has emerged in recent years as a highly effective paradigm for understanding how the brain functions (Clark, 2013; Friston, 2018). It was applied initially to sensory processing, including the visual system (Rao and Ballard, 1999), and has since been extended to include cognition and emotion (Barrett, 2017). This framework can also be used to understand psychiatric disorders such as depression and other stress-related disorders (Kaye and Krystal, 2020; Smith et al., 2021; Wilkinson et al., 2017). Given the broad relevance of predictive coding, a knowledge of the anatomical and physiological mechanisms underlying its implementation is of considerable interest. A key feature of predictive coding is the updating of expectations based on experience, and this involves the release of neuromodulators that indicate the sign and magnitude of error, as well as precision of prediction. The habenula is one regulator of these modulators, and the pattern of evoked activity and effects of manipulation are consistent with a role in predictive coding underlying motivated behaviour and response to stressors. Recent work has identified a number of mechanisms, such as regulation of membrane potential by astroglial cells (Cui et al., 2018), that influence habenula function by altering the level of spontaneous activity. Spontaneous activity determines the output of a network (Arieli et al., 1996) and is an important element of predictive coding (Hartmann et al., 2015). Here, we propose that the molecular clock, which influences spontaneous activity (Harvey et al., 2020), contributes to habenula function in predictive coding.

We have found that habenula clock in zebrafish is transcriptionally complex, consisting of multiple paralogs. Given the potential redundancy provided by this, we used expression of a truncated *clocka* gene (Dekens and Whitmore, 2008) to investigate the function of the habenula clock. This strategy is based on the discovery in mice that a truncated *clock* gene disrupts the circadian clock (Vitaterna et al., 1994), whereas a null mutant does not (DeBruyne et al., 2006). Expression of this antimorphic allele in the zebrafish pineal was previously shown to disruption in the expression of clock-regulated genes and in locomotion and sleep (Livne et al., 2016), indicating that it is an effective technique. Expression in the habenula appears to affect the molecular clock, as indicated by the change in *per3* gene expression. We did not observe a complete flattening of *per3* expression across the circadian cycle, however, indicating that inhibition is partial. While consistent with previous findings (Dekens and Whitmore, 2008), this level of inhibition appears sufficient to affect habenula function, as indicated by change in day-night variation in neuromodulators that are under the control of the habenula.

We observed that the freezing behaviour evoked by the alarm pheromone is lower at night compared to the day, and this difference is reduced by expression of the truncated clock gene specifically in the habenula. We also observed that dark avoidance induced by the alarm substance is reduced by manipulation of the habenula clock. Thus, the data here are consistent with the hypothesis that the habenula clock influences expectation of danger. Clock gene expression has been reported in the habenula of other vertebrates, including mice (Salaberry et al., 2019). A broad expression of a functional clock in the mammalian habenula is consistent with electrical recordings obtained from slices, which show a change in firing rate and resting membrane potential across the circadian cycle in both medial and lateral habenula (Sakhi et al., 2014a, 2014b; Zhao and Rusak, 2005). The ability of an intrinsic clock to influence habenula-dependent expectations may thus be conserved across vertebrates. This represents a distinct mechanism from the role of clocks in other regions in the regulation of stress (Koch et al., 2017). One implication of the findings here is that identifying clock-regulated genes that regulate spontaneous activity in the habenula may be relevant in the treatment of stress related disorders.

## Acknowledgements

We thank Yoav Gothilf for providing the pT2-UAS:EGFP-2A-ΔCLK-5xMYC construct and Marnie Halpern for providing the *TgBAC(gng8:GAL4)^c426^* line. We also thank Hugh Piggins, David Lyons, Caroline Wee and Ajay Mathuru for comments and discussion, and Erik Meijering for FeatureJ. This work was funded by grants from the Singapore Ministry of Education under its Academic Research Fund Tier 2 (MOE2017-T2-058) and the National Research Foundation (NRF2017-NRF-ISF002-2676) to SJ, an ARAP fellowship from A*Star to AB, and a Start-Up Grant from the Lee Kong Chian School of Medicine to SRL. The funding agencies played no role in the study design or decision to publish.

## Author contributions

AB, RKC, JCSM, GTJH and SJ performed experiments. AB, RKC, JCSM, S, GD, SL and SJ analysed data. AB and SJ designed the study and wrote the manuscript.

---

**Figure S1.**
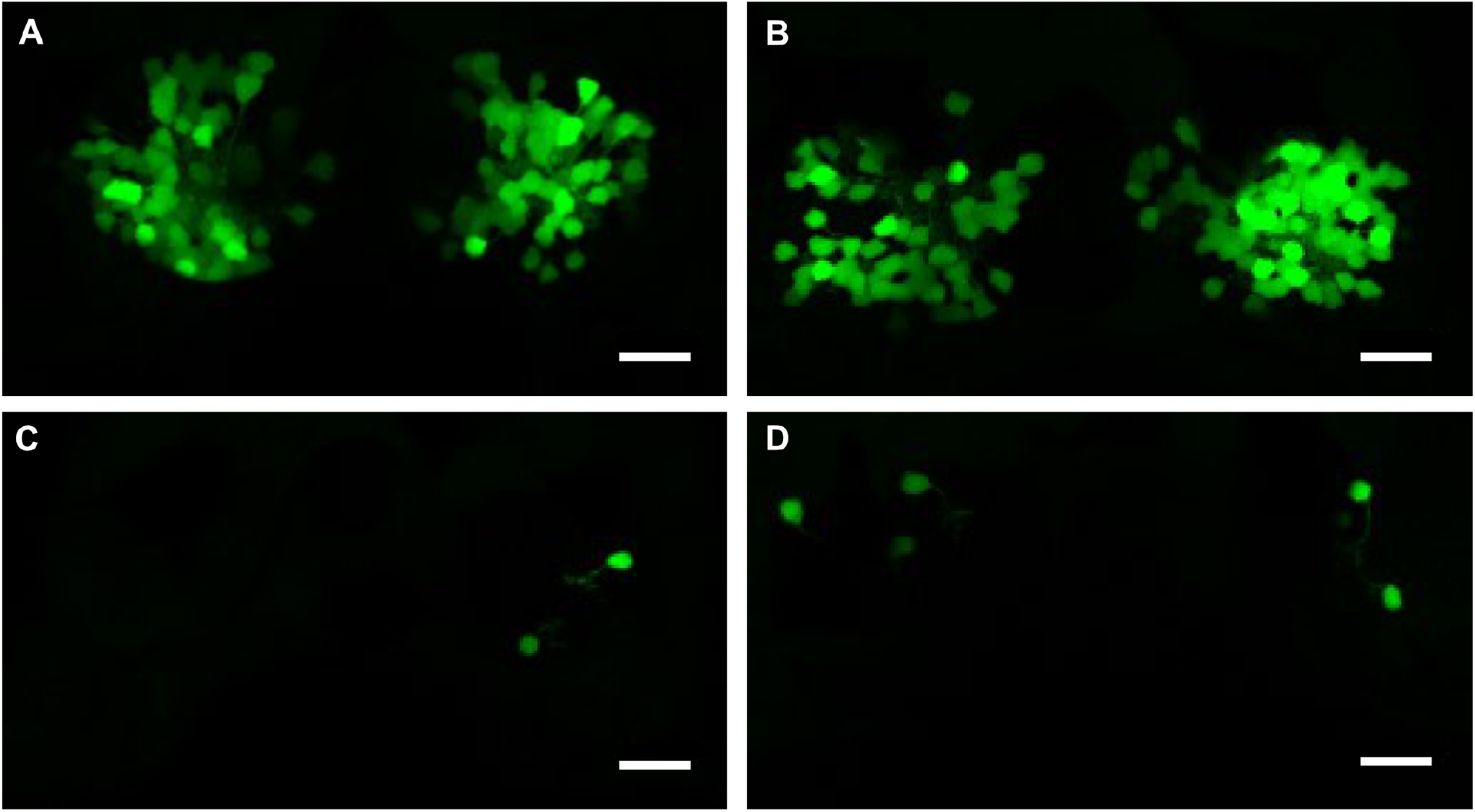
Examples of *Tg(gng8:GAL4, UAS:EGFP-*Δ*clocka)* larval zebrafish with broad (A, B) and sparse (C, D) expression of EGFP in the habenula. All images are dorsal view, with anterior to the top. Scale bar = 10 μm.

